# Differential glycosylation of alpha-1-acid glycoprotein (AGP-1) contributes to its functional diversity

**DOI:** 10.1101/2020.02.27.968974

**Authors:** Mosale Seetharam Sumanth, Shancy P Jacob, Kandahalli Venkataranganayaka Abhilasha, Bhanu Kanth Manne, Venkatesha Basrur, Sylvain Lehoux, Robert A Campbell, Christian C Yost, Matthew T Rhondina, Thomas M McIntyre, Richard D Cummings, Andrew S Weyrich, Gopal K Marathe

**Author notes:** To whom correspondence should be adressed: Gopla K Marathe, Department of Studies in Biochemistry, University of Mysore, Manasagangothri, Mysuru – 570006, Karnataka, India.

## Abstract

Alpha-1-acid glycoprotein (AGP-1) is a positive acute phase glycoprotein with uncertain functions. Serum AGP-1 (sAGP-1) is primarily derived from hepatocytes and circulates as 12 to 20 different glycoforms. We isolated a glycoform secreted from stimulated human neutrophils (nAGP-1). Its peptide sequence was identical to hepatocyte-derived sAGP-1, but nAGP-1 differed from sAGP-1 in its chromatographic behaviour, electrophoretic mobility, and glycosylation. The function of these two glycoforms also differed. sAGP-1 activated neutrophil adhesion, migration and NETosis in a dose-dependent fashion, while nAGP-1 was ineffective as an agonist for these events. Furthermore, sAGP-1, but not nAGP-1, inhibited LPS-stimulated NETosis. However, nAGP-1 inhibited sAGP-1-stimulated neutrophil NETosis. The discordant effect of the differentially glycosylated AGP-1 glycoforms was also observed in platelets where neither of the AGP-1 glycoforms alone stimulated aggregation of washed human platelets, but sAGP-1, and not nAGP-1, inhibited aggregation induced by Platelet-activating Factor (PAF) or ADP, but not by thrombin. These functional effects of sAGP-1 correlated with intracellular cAMP accumulation and were accompanied by phosphorylation of the PKA substrate Vasodialator stimulated phosphoprotein (VASP) and reduction of Akt, ERK, and p38 phosphorylation. Thus, the sAGP-1 glycoform limits platelet reactivity while nAGP-1 glycoform also limits pro-inflammatory actions of sAGP-1. These studies identify new functions for this acute phase glycoprotein and demonstrate that the glycosylation of AGP-1 controls its effects on two critical cells of acute inflammation.

## Introduction

Alpha-1-acid glycoprotein (AGP-1) is a positive acute phase glycoprotein with a low pI of 2.8 - 3.8 and a carbohydrate content contributing to 45% of its total mass. Hepatic synthesis of AGP-1 increases during an inflammatory response (1,2) and is released to the circulation (sAGP-1). Although numerous activities like immunomodulation and altering inflammatory milieu have been ascribed to AGP-1 (2,3) its function(s) is ill defined. Other cell types including human breast epithelial cells, lymphocytes and monocytes secrete AGP-1 in response to appropriate inflammatory stimuli (3–5). In addition, human neutrophils, secrete AGP-1 in response to platelet-activating factor (PAF), lipopolysaccharide (LPS), TNFα and phorbol myristyl acetate (PMA) (6). AGP-1 undergoes extensive glycosylation and approximately 12-20 different glycoforms of AGP-1 are present in human blood (2,7). This may be an underestimate, as prior studies using capillary electrophoresis-mass spectrometry, indicated that more than 150 isoforms of AGP-1 occur in human plasma (8,9).

This impressive number of glycoforms of AGP-1 is altered during acute and chronic inflammation (5,10–13). There is ample evidence to support the concept that the glycosylation pattern and degree of chain branching may serve as a marker for specific disease conditions (3,9,10,12–17). However, unravelling AGP-1 function(s) remains challenging. For example, AGP-1 offers non-specific protection against various agonists - induced platelet aggregation, Gram-negative bacterial infection and TNFα-induced lethality (18–20), but also promotes monocytes to an anti-inflammatory M2 phenotype rendering them ineffective against bacterial infections (21). Furthermore, AGP-1 inhibits neutrophil migration in sepsis (22), contributing to infection. More recently, Higuchi et al. (23) have shown that AGP-1 is also involved in the allograft rejection after kidney transplantation.

We previously observed that serum AGP-1 preferentially inhibits the TLR-4 agonist bacterial lipopolysaccharide (LPS), but not TLR-2 agonist Braun Lipoprotein (BLP) mediated inflammatory responses both *in vivo* and *in vitro* (24,25). We now show that the hepatocyte-derived serum AGP-1 (sAGP-1) is pro-inflammatory, while the neutrophil derived glycoforms (nAGP-1) primarily display anti-inflammatory activity. As the protein sequences of these two glycoforms are identical, our study demonstrates that carbohydrate diversity modulates AGP-1 function and contributes to its contradictory roles in inflammatory milieu.

## Results

### Neutrophils secrete distinct AGP-1 isoforms

To confirm the extra-hepatic secretion of AGP-1, freshly isolated human neutrophils (1 × 10^6^ cells/ml) were stimulated with the agonists PAF (10^−6^ M), TNFα (1000 U/ml), PMA (5 μg/ml) or LPS (1 μg/ml) and the secreted proteins were immunoblotted against AGP-1. Quiescent neutrophils did not secrete AGP-1, while neutrophils stimulated by these agonists did, suggesting the extra-hepatic source of AGP-1 (Fig. 1A). Stimulated neutrophils, however, secreted both a 43 kDa AGP-1 corresponding to sAGP-1, but also released more slowly migrating (~60 kDa) larger form of AGP-1 which we termed nAGP-1. The ratio of the two isoforms differed according to the inciting agonists. We sought to purify the higher molecular weight nAGP-1 isoform using conventional chromatographic techniques (Methods) (24) and so stimulated neutrophils with PAF to obtain an AGP-1 corresponding to sAGP-1 and nAGP-1. However, only the larger nAGP-1 was isolated by this purification protocol and was used for further experimentations (Fig. 1B).

**Fig. 1:**
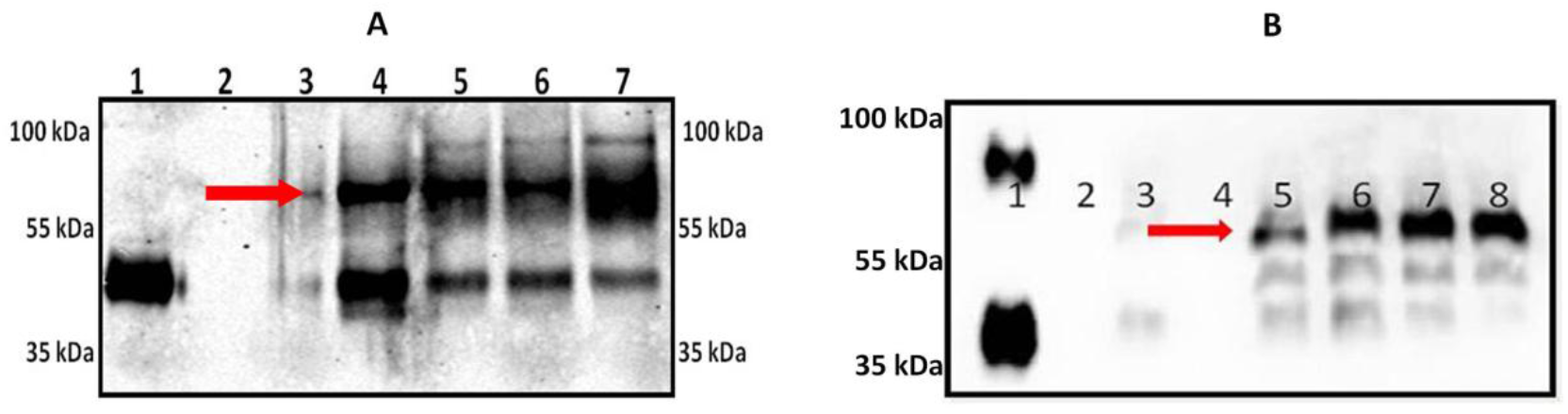
Stimulated human neutrophils secretes AGP-1 that differes from sAGP-1: **(A)** Freshly isolated human neutrophils secrete AGP-1 after stimulation (for 60 min). Secreted proteins were concentrated, resolved, and immunoblotted against AGP-1. Lanes represent; 1: sAGP-1 (250 ng); 2: 0 min Control; 3: 60 min Control; 4: PAF (10^−6^ M); 5: TNF-α (1000U/ml); 6: LPS (1μg/ml); 7: PMA (5μg/ml). **(B)** Secretion of nAGP-1 by PAF stimulated neutrophils; Lane 1: commercial AGP-1 (250 ng); 2: 0 min Control; 3: 60 min Control; 4: empty well; 5: PAF (10^−4^ M); 6: PAF (10^−6^ M); 7: PAF (10^−8^ M); 8: PAF (10^−10^ M). nAGP-1 is indicated by red arrow.

In the next series of experiments, we determined the basis for the distinct physical properties of the two AGP-1 isoforms. Unlike sAGP-1, nAGP-1 failed to bind to the DEAE-cellulose anion exchange column and so eluted in the void volume of this chromatographic separation (Figs. 2 A–D). The higher apparent molecular weight (~90 kDa) (Fig. 1B) band of commercially prepared AGP-1 suggests that either AGP-1 aggregates or this band represents additional glycoforms of AGP-1. The endotoxin content of the purified nAGP-1 was < 5 EU/mg protein, and so would not be relevant in subsequent biological analyses.

**Fig. 2:**
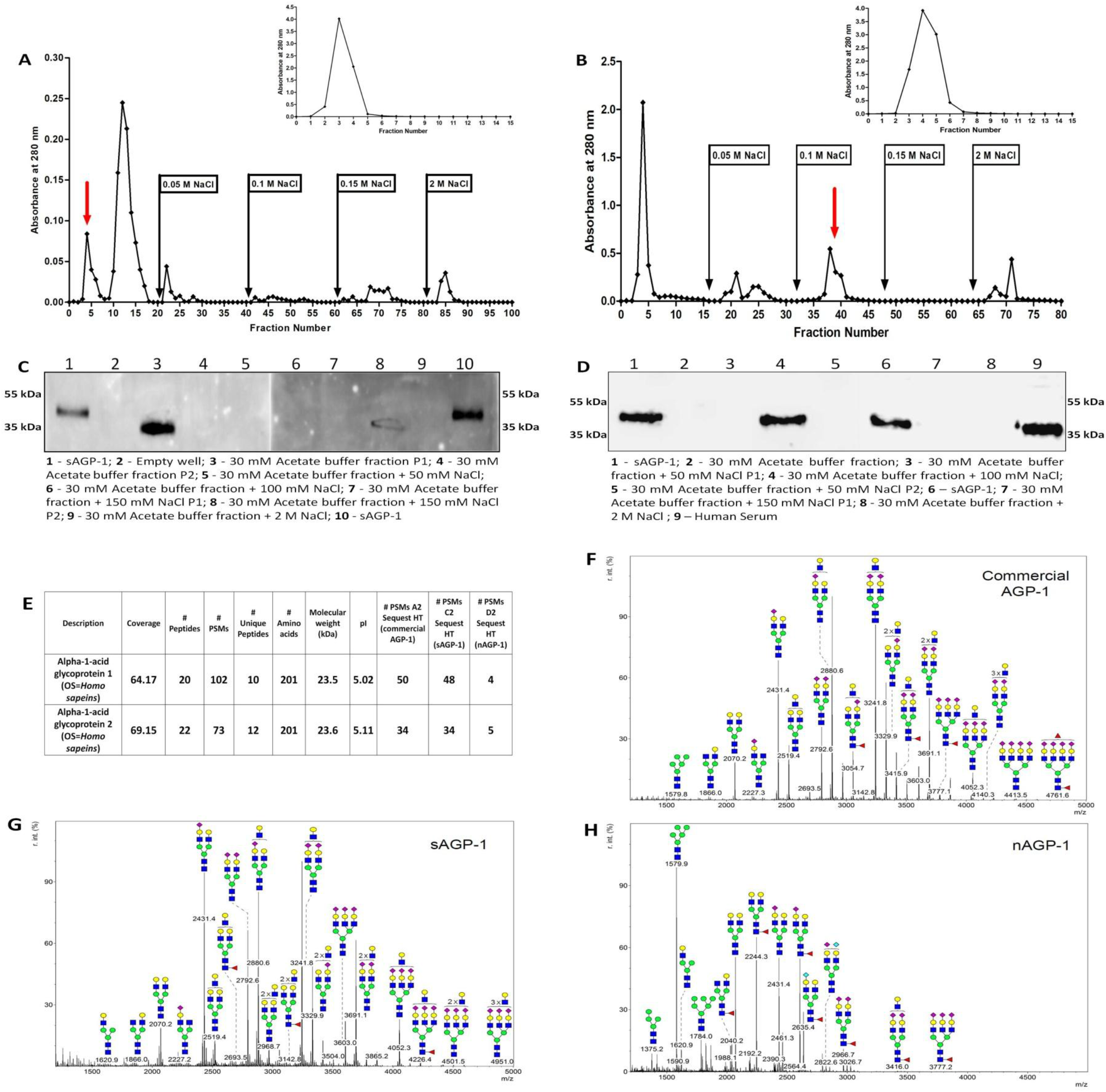
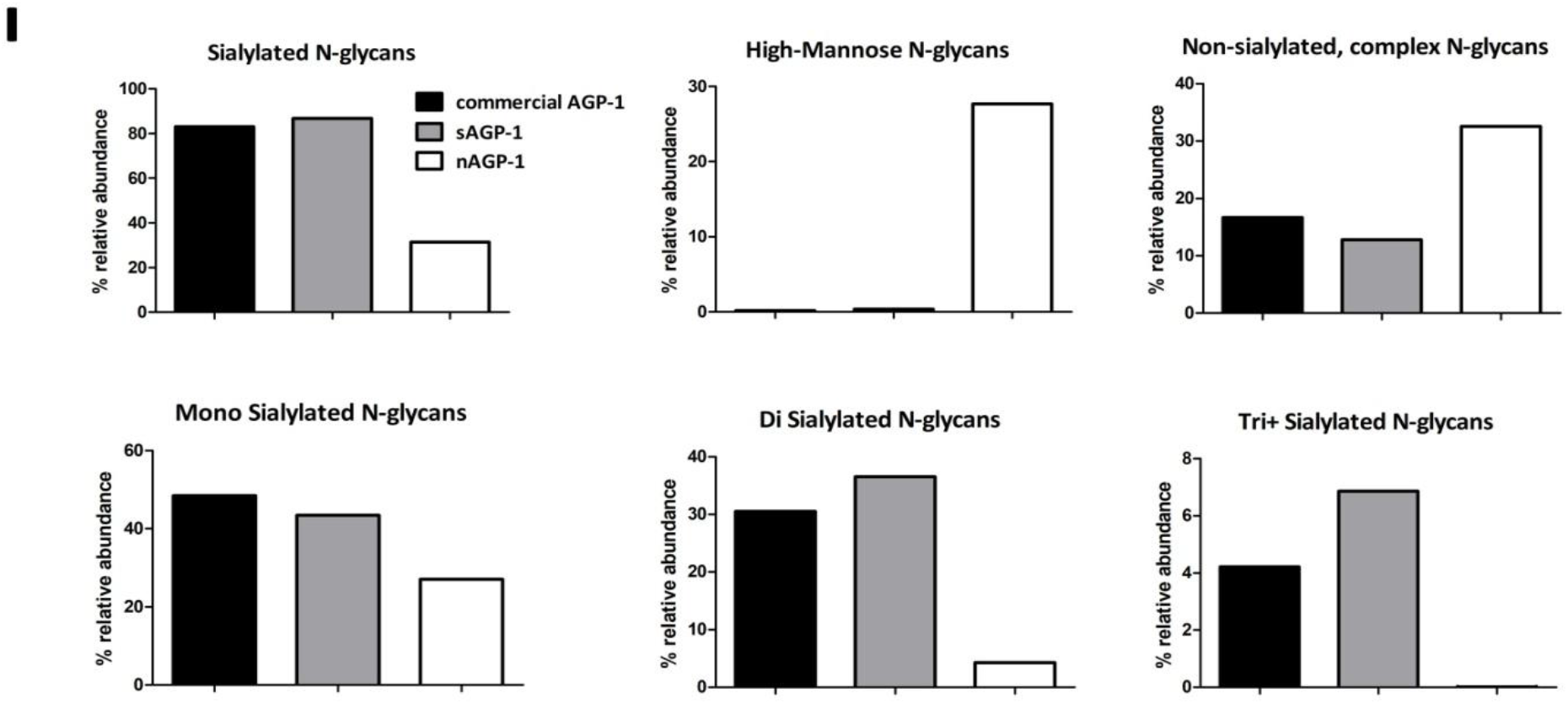
Purification and Characterization of nAGP-1 in comparison with sAGP-1: Elution profile of **(A)** PAF-induced Neutrophil supernatant (nAGP-1) and **(B)** pooled serum on Cibacron blue column (Inset) and concentrated void volume fraction of Cibacron blue column on DEAE-Cellulose column. The fraction(s) containing the respective AGP-1 is indicated by red arrow. **(C)** and **(D)** Western blotting analysis of different fractions of DEAE-cellulose column against nAGP-1 (two different gels) and sAGP-1 respectively. **(E)** Mass spectral analysis of commercial AGP-1, sAGP-1 and nAGP-1. The data revealed that the protein components of the two AGP-1s are identical. Glycomic analysis of **(F)** commercial AGP-1 and **(G)** sAGP-1 showed that these two preparations are similar. **(H)** nAGP-1 differed in glycosylation complexity and composition when compared to sAGP1. 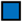 – N-Acetyl Glucoseamine; 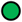 - Mannose; 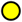 - Galactose; 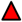 - Fucose; 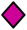 - Sialic acid. **(I)** % relative abundance of sialylated, non-sialylated, high-mannose, mono- di- and tri+ - sialylated N-glycans present in commercial AGP-1, sAGP-1 and nAGP-1.

### Glycan analysis of nAGP-1 by mass spectrometry

The electrophoretic mobility of purified nAGP-1 was slower than that of sAGP-1. To understand whether this reflects size or charge differences, we subjected the two isoforms to mass spectrometry analysis. We found the sequence of the two proteins aligned and generated peptides that were identical for the entire sequence (Fig. 2E). We next examined the glycosylation, more precisely the N-linked glycans (N-glycans), of sAGP-1 and nAGP-1. sAGP-1 is mainly glycosylated with complex bi- and tri-antennary, sialylated N-glycan chains. 86% of sAGP-1 N-glycans are sialylated, with the majority of them being mono- and di-sialylated N-glycans (Figs. 2G, I). This is similar, if not identical, to commercially available, serum-derived AGP-1 (Figs. 2F, I). By contrast, nAGP-1 has different types of N-glycan. The relative abundance of complex sialylated N-glycans in nAGP-1 was significantly reduced (~31%), and these are mainly mono-sialylated N-glycans(Figs. 2H, I). High-mannose-type N-glycans are, however, much more relatively abundant among nAGP-1 N-glycans (~27%) compared to sAGP-1 and commercial AGP-1 N-glycans (<1% for both) (Figs. 2F–I). Since nAGP-1 differs from the sAGP-1 only with respect to N-glycans and not the protein core, we considered these as distinct glycoforms of AGP-1.

### AGP-1 glycoforms differentially stimulate neutrophil adhesion

AGP-1 is elaborated in response to systemic inflammation, but with incompletely defined biological effects. To determine whether either or both of the AGP-1 glycoforms contribute to the inflammatory responses, we determined whether their undefined functions include the ability to localize neutrophils to sites of inflammation. Neutrophils rapidly and avidly adhere in response to inflammatory stimuli (26,27). To quantify this, we labelled neutrophils with Calcein, a fluorescent vital dye and assessed adhesion to a gelatin-coated glass surface that prevents interaction with unstimulated neutrophils. We found both sAGP-1 and nAGP-1 were agonists for this event, with sAGP-1 ultimately stimulating twice the number of neutrophils to adhere than nAGP-1 (Figs. 3 A, B). At lower concentrations, sAGP-1 also proved to be a significantly more potent agonist than nAGP-1. We next examined neutrophil chemotaxis using Transwell chambers to find migration in response to AGP-1 recapitulated adhesion, with sAGP-1 being significantly more potent and more stimulatory than nAGP-1 (Fig. 3C). Thus while both glycoforms of AGP-1 are neutrophil agonists, quantitative examination showed that these glycoforms differentially affect neutrophil function, with sAGP-1 acting as a more robust neutrophil chemo-attractant.

**Fig. 3:**
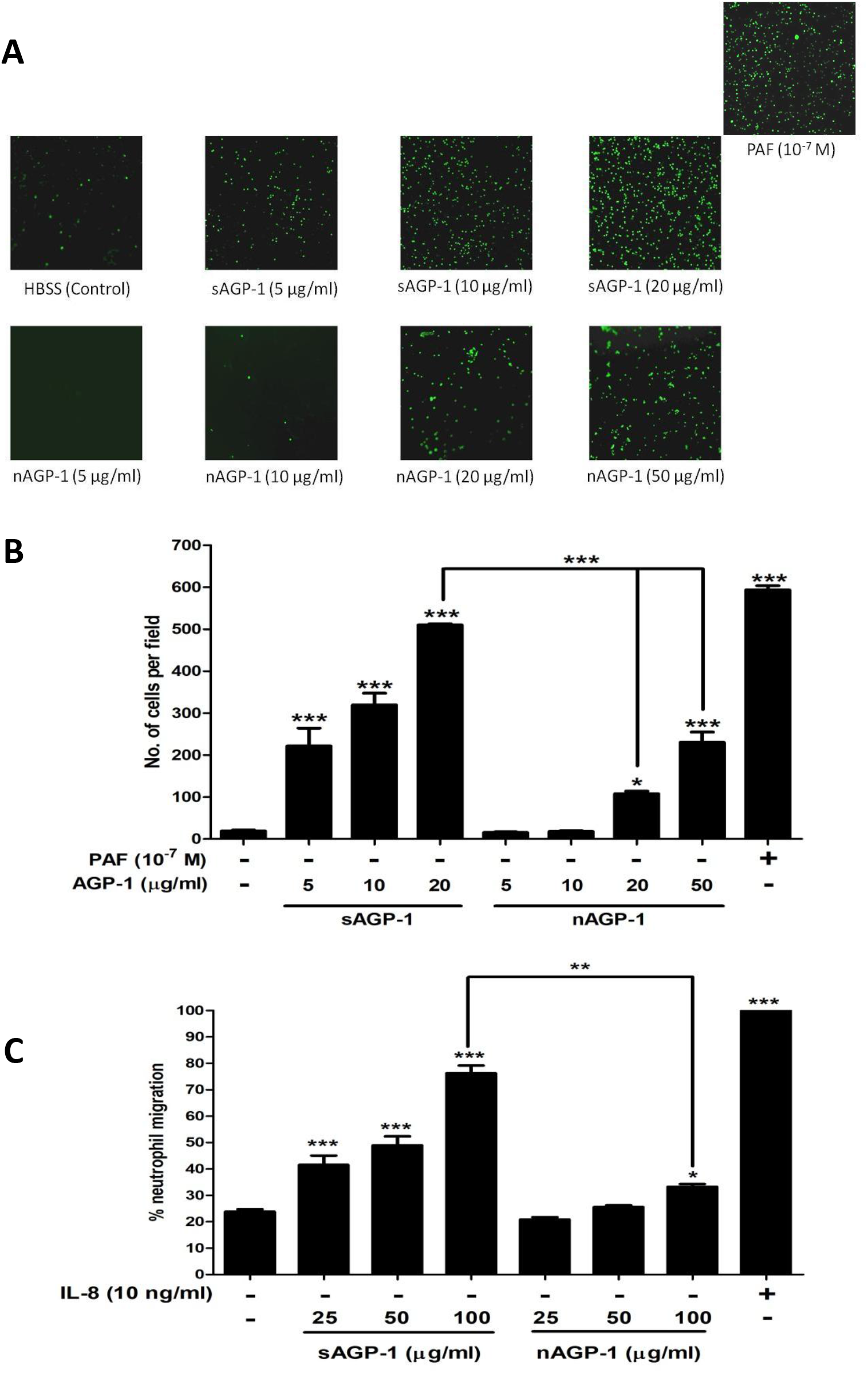
sAGP-1 is more potent activator of human neutrophils than nAGP-1 : **(A)** Adhesion. PMNs loaded with Calcein-AM in HBSS/A were treated with increasing concentrations of sAGP-1 and nAGP-1(5–50 μg/ml) separately. The incubation continued for 60 minutes at 37 °C and the non-adherent PMNs were removed by washing with HBSS. The adherent PMNs were visualized under fluorescent microscope at a magnification of 10X. PAF (10^−7^ M) was used as positive control. **(B)** AGP-1 induced PMN activation was quantified by counting the cells per field using ImageJ software as explained under ‘Methods’, and the data shown are mean ± SEM (n = 3). ***=P <0.0001, *=P<0.01. **(C)** Chemotaxis in response to sAGP-1 and nAGP-1 (25, 50 and 100 μg/ml) was tested using Transwell plates with 5μm pore size inserts (Boyden chamber assay). IL-8 (10 ng/ml) served as the positive control. Percentage of neutrophils migrated to the lower chamber were calculated and plotted. Chemotaxis induced in response to IL-8 was considered 100%. The data shown are mean ± SEM (n = 3). ***=P <0.0001, **=P<0.001 and *=P<0.01.

### sAGP-1 induces neutrophil extracellular traps (NETosis) while nAGP-1 is inhibitory

Neutrophils extrude their DNA along with antimicrobial enzymes and peptides after encountering microbes or during inflammation that forms a net-like structure to entrap bacteria in a process termed NETosis (28). We determined whether AGP-1s activate neutrophils to undergo NETosis to find sAGP-1, but not nAGP-1, induced this response (Fig. 4A). In fact, the highest concentration of sAGP-1 was comparable to the level of NETosis induced by the positive control LPS (Fig. 4B). The concentration-response relationship showed 50 μg/ml sAGP-1 was suboptimal, enabling us to determine the combined effect of sAGP-1 and nAGP-1 at this concentration. We found nAGP-1 at this concentration not only failed to induce NETosis, it inhibited the NETosis induced by sAGP-1 (Figs. 4 A, B inset). sAGP-1 suppressed LPS-induced inflammatory responses (24), and sAGP-1, but not nAGP-1, inhibited LPS-induced NETosis in a concentration-dependent fashion (Fig. 5). nAGP-1, in contrast, failed to reduce LPS-induced NETosis even at 100 μg/ml, where sAGP-1 abolished the NETosis induced by LPS (Fig. 5).

**Fig. 4:**
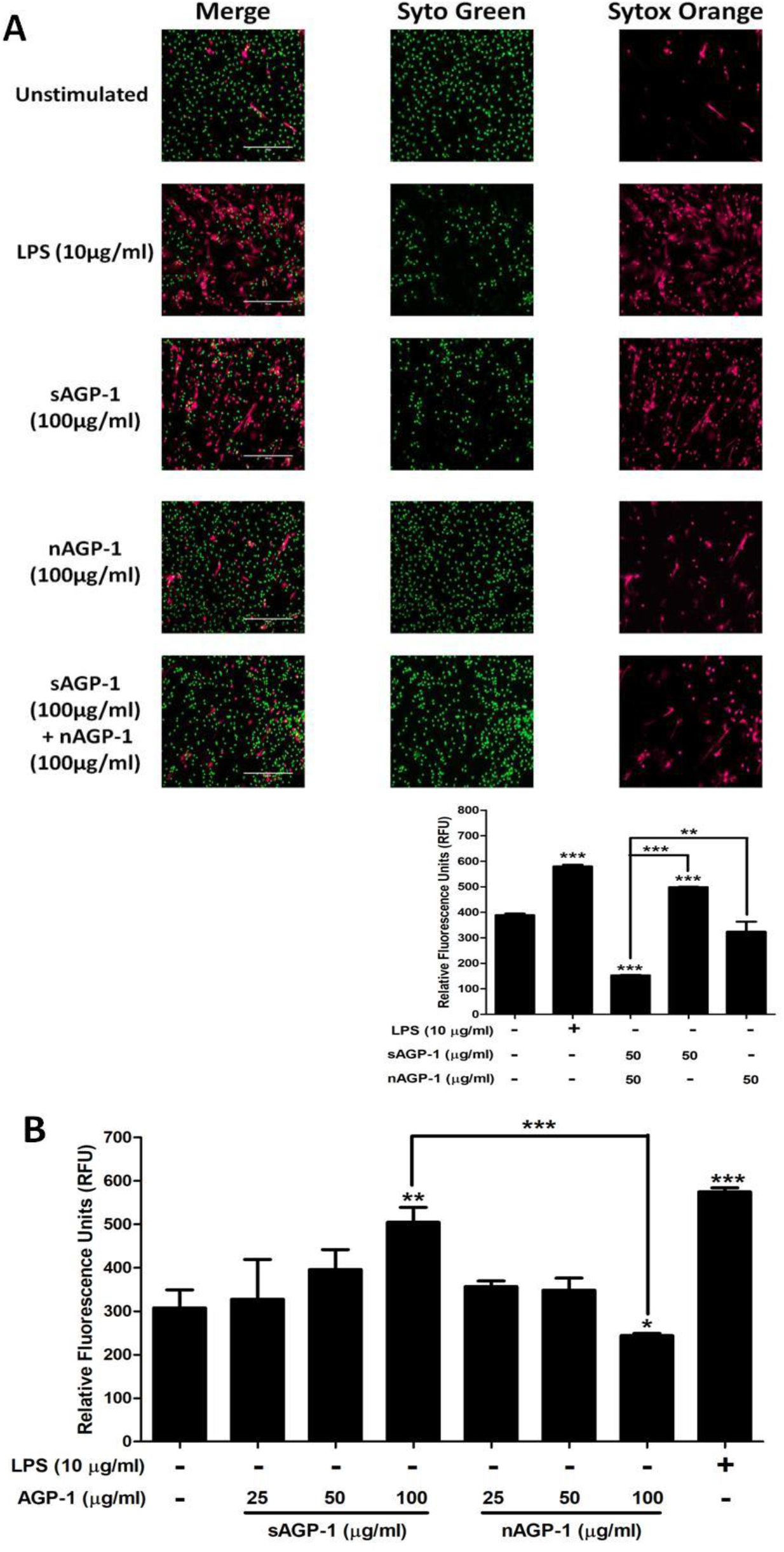
sAGP-1, but not nAGP-1, induces NETosis: **(A)** NET formation. Isolated human neutrophils adhering to poly-L-Lysine coated slides were incubated with media alone, or with sAGP-1, nAGP-1 and LPS for 1 hour and were then stained with cell-permeable, Syto Green, and/ or with cell-impermeable Sytox Orange to quantify cells lacking a permeability barrier, to label polymeric DNA. NETs were assessed by live cell imaging using fluorescence microscope at 20× magnification (Scale: 200μm). NET formation by LPS (red fluorescence) served as positive control. **(B)** Concentration-response relationships. NETs were quantified using high-throughput method as explained in the “Methods” section. Neutrophils were stimulated with sAGP-1, nAGP-1 (25, 50 and 100 μg/ml) or LPS for 1 hour before quantifying NETs by fluorometry. In a parallel experiment, neutrophils were incubated with sAGP-1, nAGP-1 and combination of sAGP-1 and nAGP-1 (inset). LPS served as positive control. The data shown are mean ± SEM (n = 3). ***=P <0.0001, **=P<0.001 and *=P<0.01.

**Fig. 5:**
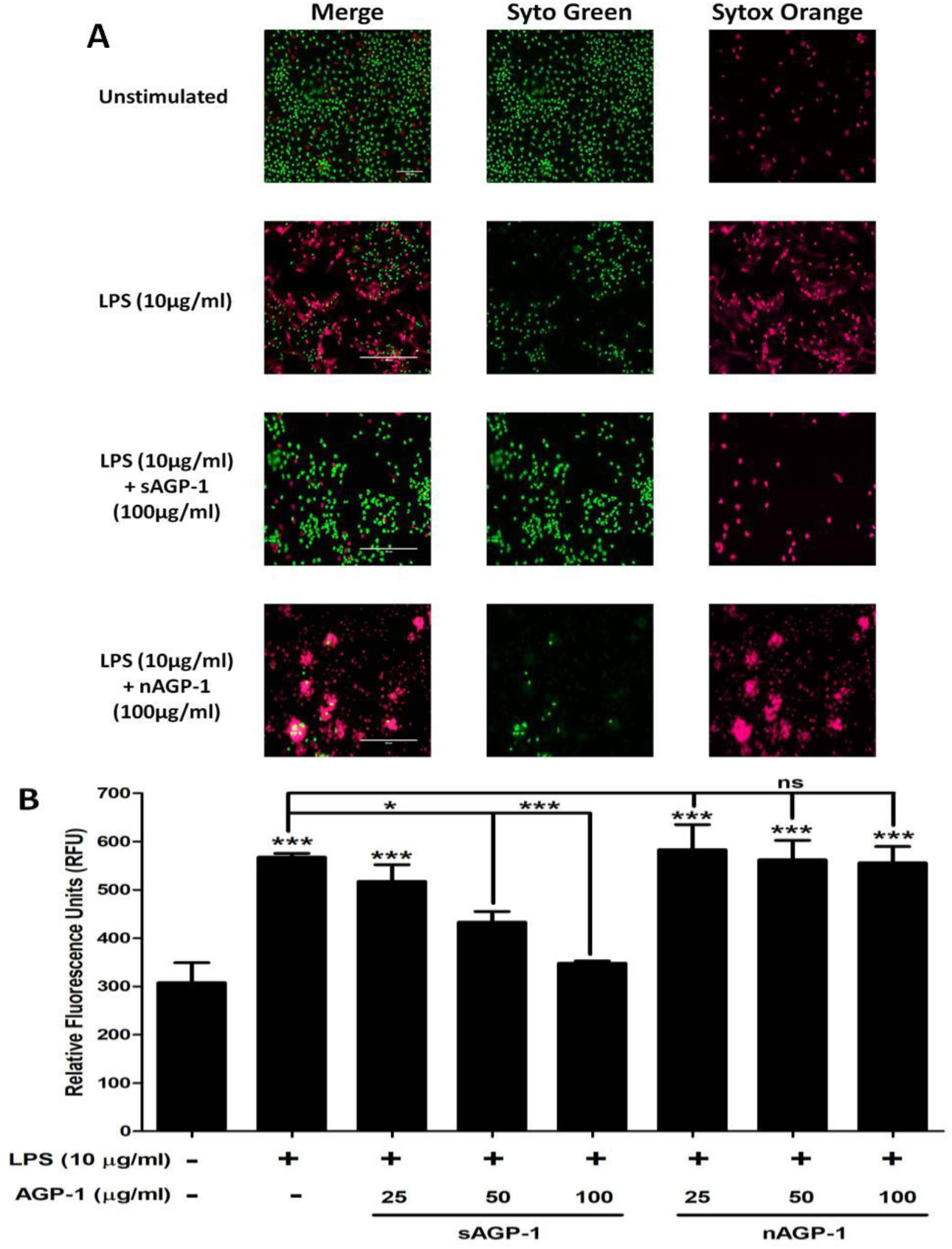
sAGP-1 inhibits LPS-induced NETosis: **(A)** Neutrophils were treated respectively with vehicle, LPS (10 μg/ml) in the presence/absence of sAGP-1 and nAGP-1 (25, 50 and 100 μg/ml). The mixture was incubated for 60 min at 37 °C and stained with Syto Green and Sytox Orange fluorescent dye mixture. The NETs were visualized under a fluorescent microscope at a magnification of 20×. sAGP-1 inhibited TLR-4 (LPS) mediated NETosis, while nAGP-1 did not have any effect on LPS – mediated effects. **(B)** NETosis was quantified using high-throughput method as explained under “Methods”. The data shown are mean ± SEM (n = 3). ***p<0.0001, *p<0.01 as determined by ANOVA.

### sAGP-1, but not nAGP-1, suppresses platelet aggregation induced by incomplete stimuli

Platelets are critical players in inflammation (29). To understand whether AGP-1 affects these cells, we examined aggregation of washed human platelets incubated with AGP-1 alone or together with inflammatory agonists. We found neither of the AGP-1 glycoforms stimulated platelet aggregation by themselves (Fig. 6A inset). Instead, sAGP-1 profoundly suppressed, and ultimately abolished, platelet aggregation induced by either PAF (Figs. 6A, E & F) or ADP (Figs. 6B, G & H). In contrast, nAGP-1 was without effect on stimulated platelet aggregation. Platelet aggregation in response to the single “strong” platelet agonist thrombin, however, was not affected by either of the AGP-1 glycoforms (Figs. 6C, I & J), while soluble collagen acting on non-G protein-coupled receptors was only modestly affected (Figs. 6D, K & L).

**Fig. 6:**
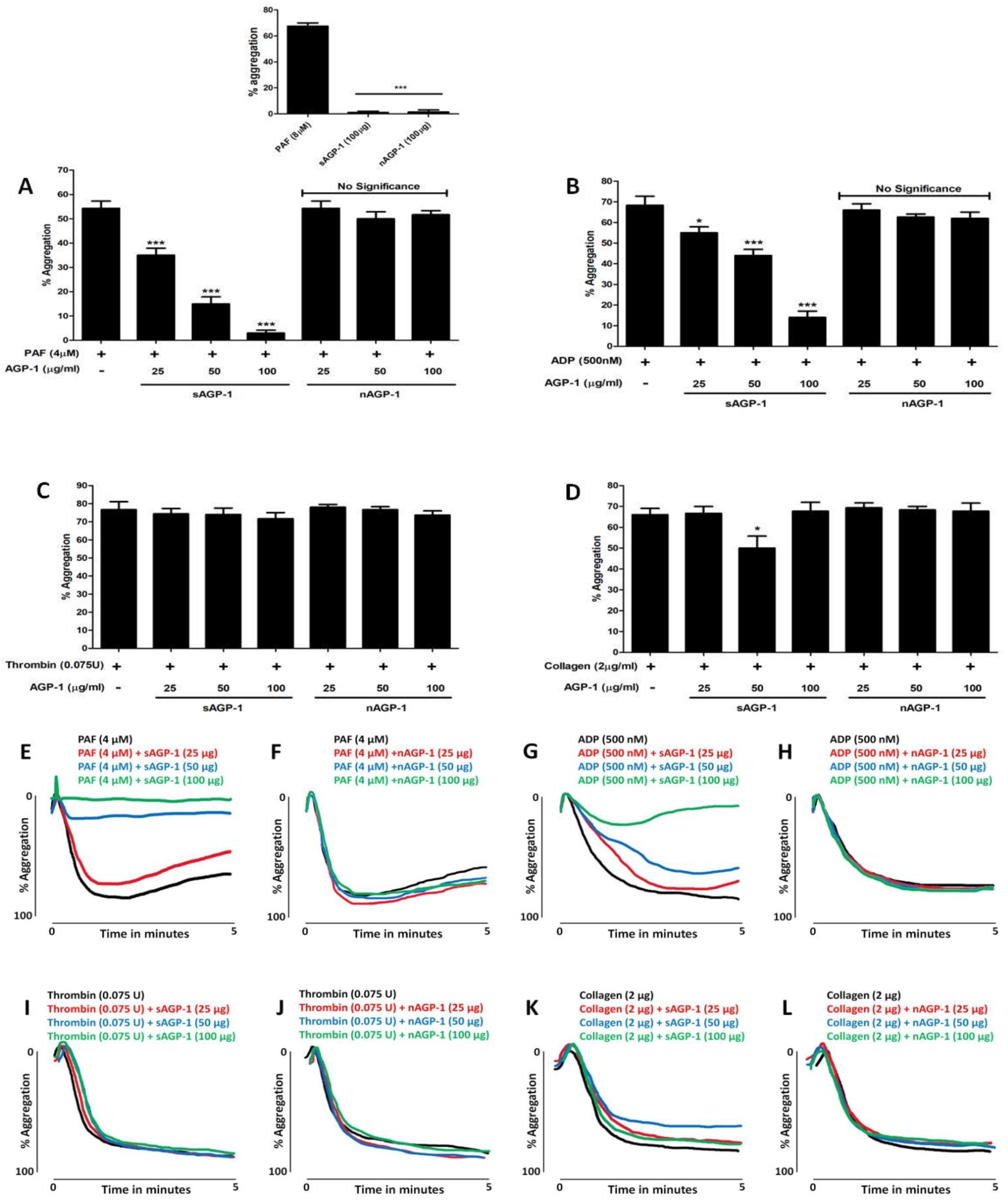
sAGP-1, but not nAGP-1, inhibits stimulated platelet aggregation. Platelet aggregation was induced by indicated amounts of PAF **(A, E & F)**, ADP **(B, G & H)**, thrombin **(C, I & J)**or collagen **(D, K & L)** with or without sAGP-1 and nAGP-1. In a parallel experiment, platelet aggregation was induced by the stated amount of sAGP-1 and nAGP-1 using PAF – induced aggregation as positive control **(A inset)**. Each set of experiments wer carried out with platelets from the same donor as well as repeated with platelets from at least three other donors. All assays were performed using Chrono-log aggregometer and the traces were recorded using AGROLINK software. Representative trace of three independent experiments is shown here. The aggregation traces were merged using Wacom graphics pad. The data shown are mean ± SEM (n = 3). ***p<0.0001, *p<0.01 as determined by ANOVA.

### sAGP-1 stimulates cAMP accumulation in platelets stimulated by incomplete agonists

To understand the molecular mechanism underlying sAGP-1 inhibition of platelet aggregation, we assessed the effect of sAGP-1 on inhibitory cyclic AMP (cAMP). We found sAGP-1 modestly, but not significantly, reduced cAMP levels in quiescent platelets, while the strong agonist thrombin significantly reduced cAMP abundance (Fig. 7A). sAGP-1 did not affect thrombin-suppressed cAMP, but did greatly increase cAMP levels in platelets stimulated with either PAF or ADP. We next determined whether sAGP-1 affected kinase signalling, and visualized the phosphorylation status of Akt, p38, and ERK kinases as well as the actin regulator vasodilator activated phosphoprotein (VASP) that is phosphorylated by cAMP and protein kinase A (PKA). The phosphoblot of platelets stimulated in the presence or absence of sAGP-1 showed that thrombin stimulation, like aggregation, was unaffected by this acute phase protein (Figs. 7B, C–F). These experiments also demonstrated that VASP phosphorylation was increased by sAGP-1 when cAMP levels increased (Figs. 7 A, B and F). In contrast, phosphorylation and activation of serine kinases by PAF or ADP was reduced by sAGP-1 (Figs. 7 B-F). These findings were confirmed by the results of three separate experiments that used platelets from different donors. The totality of these results shows congruence of the effect of sAGP-1 on aggregation, cAMP accumulation, kinase phosphorylation, and VASP phosphorylation across a range of agonist effectiveness.

**Fig. 7:**
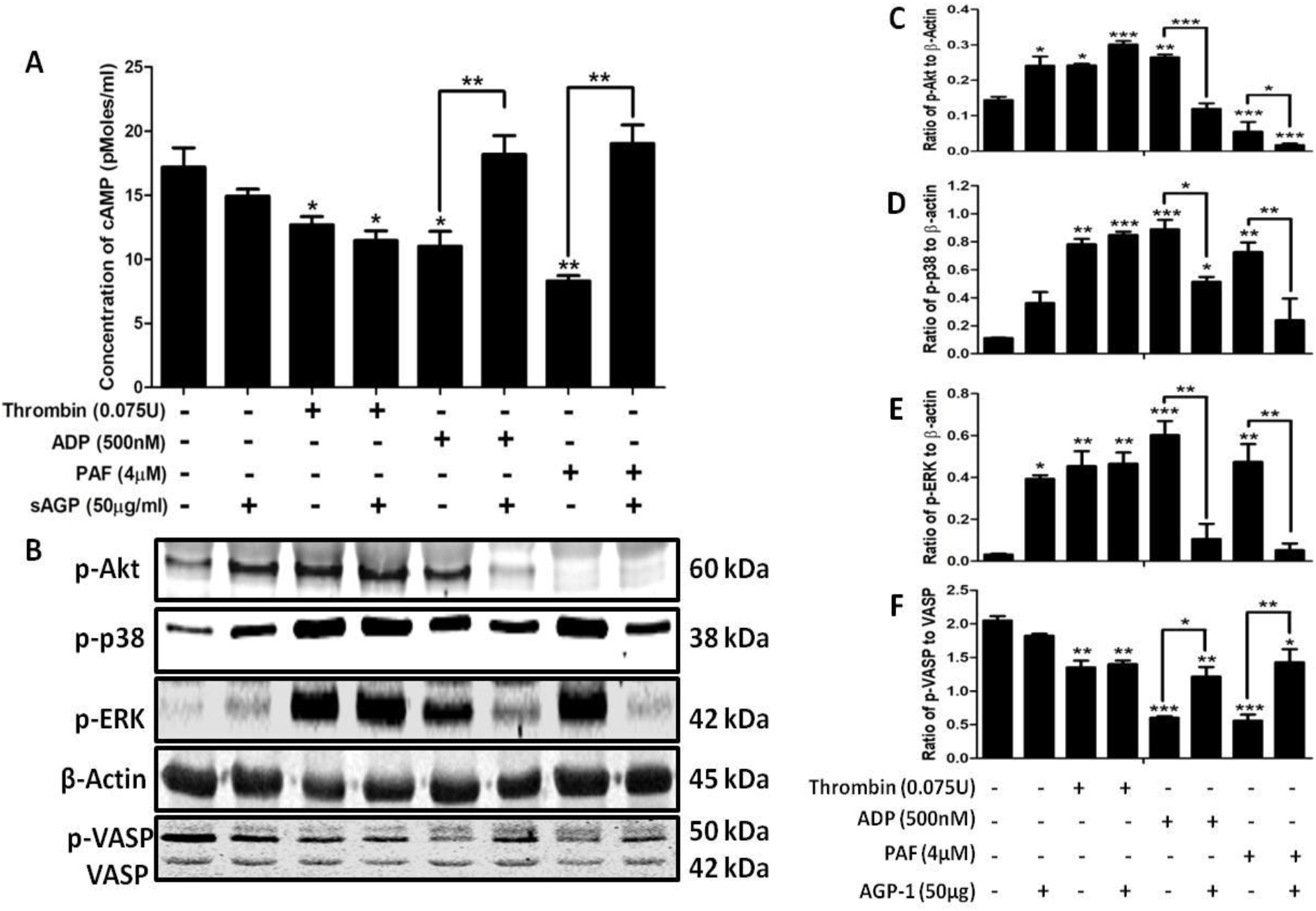
Inhibition of PAF and ADP-induced platelet aggregation by sAGP-1 is mediated by a cAMP-dependent pathway: **(A)** cAMP levels. Washed platelets were stimulated with PAF, ADP or Thrombin with or without sAGP-1. After incubation, the cells were collected by centrifugation and then assayed for cAMP by ELISA. The data represents results from two different experiments. The data shown are mean ±SEM **p < 0.01 and *p < 0.05 as analysed by one way ANOVA. **(B)** Phosphokinase western blot. The phosphorylation status of the stated kinases treated, or not, with the indicated agonist as in the preceding panel was visualized by western blotting. Phosphorylation of Akt, p38, ERK, or VASP was normalized using total β-actin. **(C – F)** Densitometric analysis of phosphokinase immunoblots. Densitometric analysis of blots obtained from 3 experiments was done using ImageJ software. The data shown are mean ±SEM ***p < 0.001, **p < 0.01 and *p < 0.05.

## Discussion

Our results demonstrate unique biological effects of different glycoforms of the acute phase glycoprotein AGP-1, depending on its origin in serum (sAGP-1) or from activated neutrophils (nAGP-1), which are highly relevant to the inflammatory responses. Physiologic inflammation is a defence against microorganisms, harmful stimuli, damaged cells, or irritants (30). One group of proteins marking this process are positive acute phase proteins (31,32) like AGP-1, which is often considered as the marker of inflammation (2,33). Although AGP-1 levels are elevated during inflammation, its biological function(s) remain obscure. We previously purified serum AGP-1 (sAGP-1, 43 kDa) to explore its differential effects on TLR-2 and TLR-4 mediated inflammatory responses *in vivo* and *in vitro* (24). As there are more than 150 glycoforms of AGP-1 present in the human plasma, this could add to the complexity of AGP-1 biology (2,8,13,14,17). To this end, we isolated and characterized a novel AGP-1 glycoform from PAF-stimulated human neutrophils (nAGP-1) and compared its biological functions with the abundant form, sAGP-1.

AGP-1 is an acute phase protein secreted from liver, but AGP-1 is also secreted from extra-hepatic sites (3–5). We determined whether an extra-hepatic source of AGP-1 (neutrophils) was identical to hepatic-derived sAGP-1 in structure and function, or whether the plethora of glycoforms also contributes to functional diversity. We found that quiescent human neutrophils, the first responder of the innate immune system, secreted little AGP-1, but did so in response to any of several lipid and protein agonists, or phorbol myristic acid (PMA). The AGP-1 released from stimulated neutrophils consisted primarily of two immunoreactive species with distinct electrophoretic mobility. One glycoform migrated like sAGP-1, while the major product when PAF was the stimulus was a slower migrating glycoform. There are previous reports that a higher molecular weight AGP-1 exists in the secondary granules of both human and bovine neutrophils (6,34).

We found AGP-1 secretion from neutrophils in response to PAF varied with agonist concentration, but surprisingly when purified by Cibacron chromatography and DEAE chromatography the slowly and rapidly migrating AGP-1 glycoform was completely resolved. We discovered the difference in behavior between the glycoforms was the inability of the nAGP-1 to bind positively charged DEAE resin. The differences in behavior was not due to protein itself, as mass spectrometry evidence confirmed that the sAGP-1 retained by the DEAE column generated the same peptides after trypsin digestion as that of nAGP-1, and the same AGP-1 gene was identified by the Protein Discoverer program for both proteins. The differences between sAGP-1 and nAGP-1 were found to reside in N-glycans. Analyses revealed that the glycosylation of nAGP-1 is dramatically different from that of sAGP-1, in that nAGP-1 mainly expressed high-mannose, non-sialylated and mono-sialylated N-glycans, as opposed to sAGP-1, which expressed mono- and di-sialylated N-glycans. In accordance with this, nAGP-1, which only partially expresses mono-sialylated N-glycans, failed to be retained by the positively charged DEAE resin. Thus, we concluded that the slow and rapidly migrating AGP-1 species from stimulated neutrophils are glycoforms of the same AGP-1 protein.

Our next question was whether AGP-1 glycoforms differ in function. To define this, we first defined relevant functions for AGP-1 in the inflammatory system. AGP-1-induces Ca^2+^ influx in neutrophils (35,36) to activate these cells (24). In accordance with our previous data, sAGP-1-induces concentration-dependent activation of neutrophil adhesion and migration. Here sAGP-1 is as effective as IL-8, but nAGP-1 is significantly less potent in activating either of these responses. Similarly, we found that while sAGP-1 stimulates NETosis, nAGP-1 does not. Moreover, nAGP-1 suppresses NETosis stimulated by sAGP-1, while sAGP-1, but not nAGP-1, suppresses NETosis induced by LPS. This effect might be due to direct interaction of AGP-1with LPS thereby quenching LPS’s effects as suggested by Huang et. al. (37). To check this possibility, we performed binding studies using FITC conjugated LPS with sAGP-1. However these results were not conclusive (data not shown). Additionally, the carbohydrate moiety of nAGP-1 might physically prevent recognition of the carbohydrate structures displayed by sAPG-1 and LPS. We conclude that plasma sAGP-1 stimulates leukocyte function, while extra-hepatic nAGP-1 mostly does not. This difference in biological function reflects differences in post-translational modification of these proteins by glycans. Thus, stimulated neutrophils produce an anti-inflammatory glycoform of AGP-1 that counteracts the pro-inflammatory actions of circulating sAGP-1.

The differential effects of AGP-1 glycoforms are also extended to the responses of platelets, which play a critical role in inflammation (29,38–40). While neither glycoforms alone stimulated platelet aggregation, sAGP-1 proved to be a highly effective suppressor of platelets stimulated by PAF or ADP, less so for those stimulated by soluble collagen, and was without effect on the strong agonist thrombin. Again, nAGP-1 differed in function from sAGP-1 and had no effect on stimulated platelet aggregation. The concentration-dependent inhibitory effect of sAGP-1 on platelet function induced by PAF or ADP correlated to a sharp increase in intracellular cAMP levels. In contrast, thrombin-induced aggregation was not affected by sAGP-1 and intracellular cAMP also was unaffected by this acute phase protein. Phosphorylation of VASP is dependent on cAMP and the PKA pathway it controls (41,42), and sAGP-1 upregulated phosphorylation of VASP in PAF- or ADP-stimulated platelets, but not in thrombin stimulated cells. sAGP-1 reduced PAF- and ADP-stimulated phosphorylation and activation of Akt, ERK, and p38 kinases, but again thrombin stimulation of these enzymes were unaffected by sAGP-1. These results, then, suggest that sAGP-1 inhibits platelet aggregation by weaker agonists via cAMP-PKA mediated signaling (Fig. 8).

**Fig. 8:**
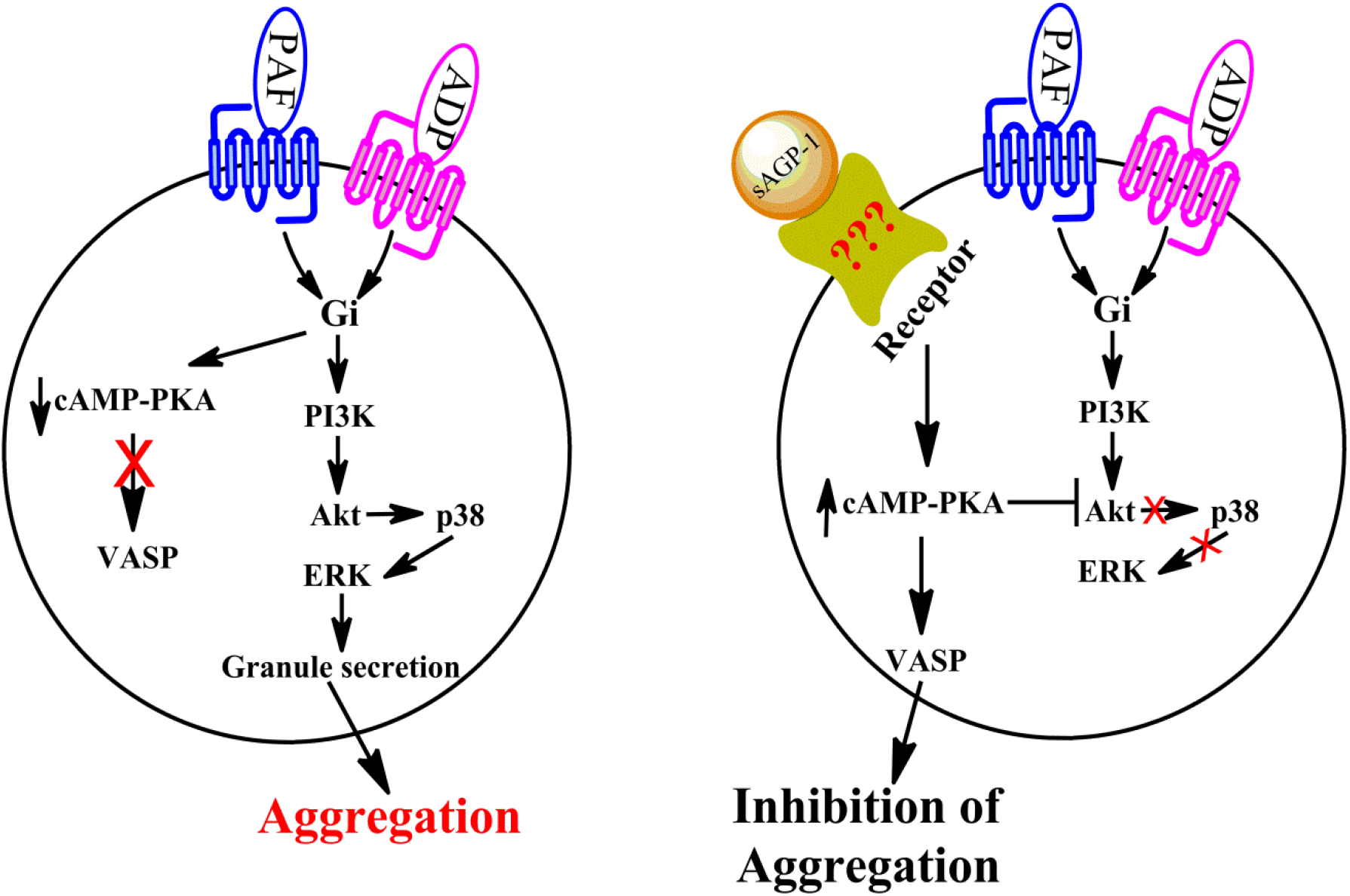
Proposed action of sAGP-1-induced inhibition of PAF- and ADP-mediated platelet aggregation: sAGP-1 up-regulates cAMP levels in PAF or ADP activated platelets, thereby activating PKA. This then activates VASP and inhibits Akt phosphorylation and activation thereby inhibiting downstream signalling and platelet aggregation.

In conclusion, our studies demonstrate that AGP-1 displays heterogeneity in its glycan structures associated with differences in physiological functions. Isolation and characterization of a glycoform of AGP-1 released from stimulated human neutrophils represents a non-hepatic source of extracellular AGP-1, which is less pro-inflammatory than hepatic sAGP-1 on platelets and neutrophils. The basis for these differences is the distinct carbohydrate chain composition and structure incorporated by the two cell types rather than the AGP-1 protein itself. Furthermore, our studies demonstrate that the AGP-1 glycoform released from stimulated neutrophils in the early phases of inflammation is not an effective positive acute phase protein, but instead counteracts the hepatic sAGP-1 glycoform. This, in turn, identifies a need for future investigations to determine functions of the multitude of other AGP-1 glycoforms and there is a need to differentially quantify AGP-1 glycoforms rather than just AGP-1 protein in health and diseases. However the greatest challenge lies in identifying appropriate receptors (Fig. 8) for AGP-1 glycoforms that ligate and alter intracellular signaling pathways.

## Materials and Methods

### Chemicals

Cibacron F3GA agarose, DEAE-cellulose gel beads, anti-human AGP-1 antibody (A5566), anti-mouse-IgG-HRP antibody, commercial AGP-1, Foetal Bovine Serum (FBS), Lipopolysaccharide (LPS), recombinant IL-8, Micrococcal DNase, Adenosine diphosphate (ADP), Thrombin, Apyrase, Prostaglandin E_1_ (PGE_1_), Medium-199, Poly-L-Lysine, Hank’s-Balanced Salt Solution (HBSS) and RPMI media were procured from Sigma Chemicals Co., St. Louis, MO, USA. Phospho-specific antibodies to p-38, JNK, ERK, VASP; β-actin, and anti-rabbit IgG-HRP were obtained from Cell Signaling Technology, Danvers, MA, USA. Complete Mini EDTA-free protease inhibitor cocktail tablets were from Roche Diagnostics, Mannheim, Germany. PVDF membrane was from BioRad Laboratories, Hercules, CA, USA and Thioglycollate media was obtained from Sisco Research Laboratories, Mumbai, India. Trypsin was purchased from Promega, Madison, WI, USA. Sytox Green, Sytox Orange, Syto Green, Calcein 2 AM and molecular weight markers were procured from Invitrogen, Carlsbad, CA, USA. Collagen was from Chronolog, Havertown, PA, USA. Platelet-activating Factor was from Avanti polar Lipids, Alabaster, AL, USA. All the reagents used in the Glycomics Massspectrometry studies were procured from Sigma Aldrich Co., St. Louis, MO, USA except for PNGaseF which was from New England Biolabs, Ipswich, MA, USA.

### Serum collection

Blood was drawn from healthy volunteers with informed consent. Permission to draw blood from the human volunteers was obtained from the Institutional Human Ethics Committee, University of Mysore, Mysuru (UOM No. 104 Ph.D/2015-16). Briefly, the blood was collected and allowed to coagulate. The coagulated blood was then centrifuged at 600 × g for 20 min at 25 °C to collect serum and stored at −20 °C till further use. All the experiments involving human blood were approved by the Institutional Human Ethics Committees of University of Mysore and University of Utah.

### Isolation of polymorphonuclear leukocytes (Neutrophils)

Neutrophils were freshly isolated by dextran sedimentation and separated by centrifugation in Ficoll density gradients (43). Neutrophil-rich pellets from this gradient was suspended in 1 ml of HBSS containing 0.2% human serum albumin (HBSS/A). For functional assays, whole blood was labelled with CD15 microbeads and the neutrophils were positively selected using AutoMACS. (Miltenyi Biotech, CA, USA)

### Secretion of AGP-1 from isolated human neutrophils

Ten million freshly isolated human neutrophils were stimulated or not with PAF (10^−6^ M), LPS (1 μg/ml), TNFα (1000 U/ml) and PMA (5 μg/ml) for 60 min in a total volume of 1ml in HBSS/A. In parallel, neutrophils isolated from the healthy volunteers were stimulated with varied concentrations of PAF (10^−4^ to 10^−10^ M) for 60 min at 37 °C. Neutrophil supernatants derived under these conditions were concentrated using Agilent spin concentrator with 10 kDa cut-off (Agilent Technologies, USA). Concentrated supernatant was immunoblotted against Anti-human AGP-1 monoclonal antibody with controls at both “0 min” and at “60 min”. sAGP-1 (250 ng) isolated in our laboratory (24) was used as the reference standard.

### Purification of AGP-1 from neutrophil supernatant

Purification of AGP-1 from PAF-stimulated neutrophil supernatants was carried out as described earlier (24). Briefly, pooled neutrophil supernatants were loaded on to a Cibacron F3GA agarose column (10 × 1.5 cm) equilibrated with 10 mM phosphate buffer (pH 7.8). Unbound proteins containing AGP-1 were eluted at a flow rate of 1 ml/min and quantified by UV absorbance at 280 nm. AGP-1 containing fractions were pooled and concentrated by spin concentrators (10 kDa cut off) (Agilent Technologies, USA). The concentrate was then applied to a DEAE-cellulose column (25 × 0.5 cm) equilibrated with 30 mM acetate buffer (pH 5.0). Fractions were eluted with a sodium chloride gradient (0 to 2 M) at a flow rate of 24 ml/ hour. AGP-1 containing fractions were pooled, concentrated, and quantified for protein using Lowry’s method (44). The endotoxin content of purified AGP-1 was assessed by Limulus amebocyte lysate (LAL) assay (Endochrome – K^TM^, Charles River, SC, USA) as per manufacturer’s instructions.

### Western blotting

AGP-1 samples were resolved on a reducing SDS-PAGE (7.5 % acrylamide) and visualized by Western blotting. Blots were stained using appropriate primary antibody (anti-human AGP-1 monoclonal antibody; 1:10000 v/v) and secondary antibody (anti-mouse IgG HRP conjugate; 1:5000 v/v). The blots were visualized using freshly prepared ECL reagent by UV-transillumination (Uvi-Tech, Cambridge, UK).

### Protein mass spectrometry analysis

Purified samples were resolved by SDS-PAGE, visualized using Coomassie stain, AGP-1 excised, and the band destained with 30% methanol for 4 hours. Upon reduction (10 mM dithiothreitol) and alkylation (65 mM 2-chloroacetamide) of the cysteines, protein was digested overnight with sequencing grade modified trypsin. The resulting peptides were resolved on a nano-capillary reverse phase column (Acclaim PepMap C18, 2 micron, 50 cm, Thermo Scientific) using a 1% acetic acid/acetonitrile gradient at 300 nl/min and directly introduced into Q Exactive HF mass spectrometer (Thermo Scientific, San Jose, CA). MS1 scans were acquired at 60 K resolution (AGC target=3e6, max IT=50ms). Data-dependent high-energy C-trap dissociation MS/MS spectra were acquired for the 20 most abundant ions (Top20) following each MS1 scan (15 K resolution; AGC target=1e5; relative CE ~28%). Proteome Discoverer (V 2.1; Thermo Scientific) software suite was used to identify the peptides by searching the HCD data against an appropriate database. Search parameters included MS1 mass tolerance of 10 ppm and fragment tolerance of 0.1 Da. False discovery rate (FDR) was determined using Fixed PSM validator and proteins/peptides with a FDR of ≤1 % were retained for further analysis.

### Glycomics mass spectrometry analysis

Coomassie-stained AGP-1 protein bands were excised and briefly washed with 400 μl of 50 mM AMBIC (ammonium bicarbonate) in 50% acetonitrile. The gel pieces were then briefly dried and 200 μl of a 10 mM DTT (1,4-Dithiothreitol) solution in 50mM AMBIC were added and incubated at 50º C for 30 mins. The DTT solution was discarded and the samples were washed with 200 μl of acetonitrile and dried. The samples were incubated with 200 μl of a 55 mM IAA (Iodoacetamide) solution in 50mM AMBIC for 30 mins at room temperature in dark. The IAA solution was then removed and the samples were washed with 500 μl of 50 mM AMBIC for 15 mins at room temperature followed by a wash with 200 μl of acetonitrile for 5 mins. The samples were dried prior to adding 500 μl of 50 mM AMBIC containing 10 μg of TPCK-treated trypsin and incubated overnight at 37º C. The tryptic digestion was terminated by boiling the sample for 3 mins and supernatants were recovered and collected in glass tubes. Further peptides were recovered by two cycles of washes with 200 μl of 50 mM AMBIC, 200 μl of 50% acetonitrile in 50 mM AMBIC and 200 μl of acetonitrile. All supernatants and washes were pooled in the same glass tube used and lyophilized. The dried peptides were resuspended in 200 μl of 50 mM AMBIC and incubated overnight with 1 μl of PNGaseF at 37º C. The enzymatic reaction was stopped by the addition of two drops of a 5% of acetic acid prior to purification of the released N-glycans over a C18 Sep-Pak (50 mg) column (Waters, Milford, MA) conditioned with 1 column volume (CV) of methanol, 1 CV of 5% of acetic acid, 1 CV of 1-propanol, and 1 CV of 5% of acetic acid. The C18 column was washed with 3 ml of 5% of acetic acid. Flow through and wash fractions were collected, pooled and lyophilized prior to permethylation. Lyophilized N-glycan samples were incubated with 1 ml of DMSO (Dimethyl Sulfoxide)-NaOH slurry solution and 500 μl of methyl iodide for 30 mins under vigorous shacking at room temperature. The reaction was stopped with 1 ml of MilliQ water and 1 ml of chloroform was added to purify out the permethylated N-glycans. The chloroform layer was washed 3 times with 3 ml of Milli-Q water and dried. The dried materials were re-dissolved in 200 μl of 50% methanol prior to be loaded into a conditioned (1 CV methanol, 1 CV MilliQ water, 1 CV acetonitrile and 1 CV Milli-Q Water) C18 Sep-Pak (50 mg) column. The C18 column was washed with 3 ml of 15% acetonitrile and then eluted with 3 ml of 50% acetonitrile. The eluted fraction was lyophilized and then redissolved in 10 μl of 75% methanol from which 1 μl was mixed with 1 μl DHB (2,5-dihydroxybenzoic acid) (5mg/ml in 50% acetonitrile with 0.1% trifluoroacetic acid) and spotted on a MALDI polished steel target plate (Bruker Daltonics, Bremen, Germany). MS data was acquired on a Bruker UltraFlex II MALDI-TOF Mass Spectrometer instrument. Reflective positive mode was used and data recorded between 1000 m/z and 5000 m/z. For each MS N-glycan profiles the aggregation of 20,000 laser shots or more were considered for data extraction. Only MS signals matching an N-glycan composition were considered for further analysis. Subsequent MS post-data acquisition analysis were made using mMass (45).

### Human neutrophil adhesion

For assessment of adhesion, the human neutrophil suspension (1×10^7^ cells/ml) was loaded with Calcein-AM to a final concentration of 1 μM prior to incubation for 45 min at 37 °C. The labelled neutrophils (1×10^6^ cells/well) were incubated with sAGP-1 or nAGP-1 (25, 50 and 100 μg/ml) in triplicate wells in twelve well cell culture plates (Nest Biotechnology Co. Ltd., China) pre-coated with 0.2% gelatin. Unbound neutrophils were removed by washing twice with HBSS/A and the adherent neutrophils were visualized and photographed at a magnification of 10× by fluorescent microscopy (Motic BA410 fluorescence microscope, Hong Kong; with Nikon DS-Qi2 camera, Japan) (25). The number of cells adhered in each well was determined by counting the cells adhered in 10 randomly chosen fields using ImageJ software and then calculating the average number of cells adhered per field.

### Neutrophil migration

Neutrophil migration was assessed using Transwell plates (Costar, Croning Inc., USA) with 5μm pore size inserts. 1×10^6^ (200μl) freshly isolated neutrophils in M-199 medium were incubated in the upper chamber of the transwell and the chemoattractant, IL-8 (10 ng/ml) and/ or AGP-1 (serum and neutrophil glycoforms) (25, 50 and 100 μg/ml) were introduced in the lower chamber. The number of neutrophils migrating to the lower chamber after incubation for 1 hour at 37° C with 5% CO_2_ were counted with a hemocytometer and expressed as % neutrophil migrated using IL-8 as positive control (100% migration).

### Neutrophil Extracellular Traps (NETosis)

1 million freshly isolated human neutrophils were treated with LPS (10 μg/ml) and/ or then with AGP-1 (serum and neutrophil glycoforms at 25, 50 and 100 μg/ml) before being transferred to Poly-L-Lysine coated coverslips. The neutrophils were incubated at 37 °C in 5% CO_2_ for 1 hour before adding Syto Green (cell-permeable) and Sytox orange (cell-impermeable) fluorescent dye mixture and visualized by fluorescent microscopy (EVOS Fluorescence microscope, Thermo Scientific, USA). The two dye images were merged using ImageJ software.

High-throughput NET quantification (46) was employed to quantify neutrophil NETs. For this, 24 well plates were pre-coated with poly-L-lysine before neutrophil addition followed by stimulation for 1 hour by defined agonists at 37 °C under 5% CO_2_. The extracellular traps were recovered by treating the neutrophils with Micrococcal DNase and the DNA stained with cell-impermeable Sytox Green dye. These were measured with a fluorescent plate-reader with excitation at 485 nm and emission at 530 nm.

### Platelet aggregation

Blood was drawn from healthy volunteers with informed consent. Platelet-rich plasma (PRP) was isolated from this blood using the method previously described by Zhou et al (47). Briefly, blood was drawn into citrated tubes (1/9) and centrifuged for 15 minutes at 45 × g at 25 °C to obtain PRP. The PRP was treated with PGE_1_ to stop the activation of platelets. PGE_1_ treated PRP was centrifuged at 225 × g for 20 min at 25 °C without breaking. The resulting platelet pellet was resuspended in HEPES-Tyrode’s buffer containing 0.02 U Apyrase to provide washed platelets. Platelet aggregation was performed by using 2×10^8^ platelets/mL in a final reaction volume of 500 μL at 37 °C with stirring at 1,200 rpm.

### Signaling by stimulated platelets

Washed platelets (2×10^8^ cells/ml) were stimulated for 5 min with thrombin (0.075 U/assay), PAF (4 μM), or ADP (500 nM) with or without sAGP-1 (50 μg/ml). Unstimulated washed platelets served as the negative control. Lysates were prepared by using perchloric acid (6 N) and reducing sample buffer (modified Laemmli buffer). Immunoblots were developed using specific primary and appropriate secondary antibodies for phospho-p38, phospho-Akt, phospho-ERK, phospho-VASP and β-actin (1:1000 v/v).

### ELISA for platelet cAMP

cAMP levels in washed platelets were quantified after treatment with buffer, Thrombin (0.075U/assay), ADP (500 nM), or PAF (4 μM) with or without sAGP-1 using competitive ELISA kit (EMSCAMPL, Invitrogen, Vienna, Austria). This system displays a detection sensitivity of 0.39 pmol/ml of cAMP according to the manufacturer’s instructions.

### Statistical analysis

All the experiments are representative of two or more independent experiments. The blots were analysed using Un-paired two-tailed t-test to compare the mean for each treatment group with the mean of the control group. All other results were analyzed using one-way analysis of variance (ANOVA) wherever applicable. All the statistical analysis was carried out using GraphPad Prism 5 software.

## Acknowledgement

The authors thank Mr. Neal Tolley and Mr. Mark Cody, in the University of Utah Molecular Medicine Program in Salt Lake City, Utah, USA for their technical assistance. The authors also thank Prof. Guy Zimmerman, University of Utah, Salt Lake City, USA and Dr. Stephen Prescott, OMRF, Oklahoma, USA for their timely advice during the course of this study.

## Author contributions

GKM conceived and designed experiments; MSS performed major experiments. KVA, SPJ, BKM and RAC performed other minor experiments reported in the manuscript; GKM, ASW, MTR, CCY and TMM, analyzed the results. VB conducted and analysed Mass-spectrometry; RDC and SL performed and analyzed Glycomics data. GKM, MTR, TMM, SL, RDC and MSS wrote and edited the manuscript.

## Funding Information

The authors thank University Grants Commission (UGC), India [for Basic Science Research fellowship – F4-1/2006(BSR)/7-366/2012(BSR) to MSS, National Fellowship for Higher Education (NFHE-ST) – F1-17/2015-16/NFST-2015-17-ST-KAR-3879/(SA-III/Website) to KVA] and Vision Group of Science & Technology (VGST), Government of Karnataka, India to the Dept. of Studies in Biochemistry, University of Mysore, Mysuru, India, and the National Center for Functional Glycomics Grant P41GM103694 (RDC and SL). This work was funded by the NHLBI (HL092746, HL126547), NICHD (HD93826), and NIA (AG048022), and supported in part by Merit Review Award Number I01 CX001696 from the United States (U.S.) Department of Veterans Affairs Clinical Sciences R&D (CSRD) Service. This material is the result of work supported with resources and the use of facilities at the George E. Wahlen VA Medical Center, in Salt Lake City, Utah. The content is solely the responsibility of the authors and does not necessarily represent the official views of the National Institutes of Health or the U.S. Government.

## Conflicts of Interest

The authors of this manuscript have no conflict of interest to declare.

